# ETAS®50 Attenuates SARS-CoV-2 Spike Protein-Induced IL-6 and IL-1β Production by Suppressing p44/42 MAPK and Akt Phosphorylation in Murine Primary Macrophages

**DOI:** 10.1101/2021.07.25.453717

**Authors:** Ken Shirato, Jun Takanari, Takako Kizaki

## Abstract

Excessive host inflammation following infection with severe acute respiratory syndrome coronavirus 2 (SARS-CoV-2) is associated with severity and mortality in coronavirus disease 2019 (COVID-19). We recently reported that the SARS-CoV-2 spike protein S1 subunit (S1) induces pro-inflammatory responses by activating toll-like receptor 4 (TLR4) signaling in macrophages. ETAS®50, a standardized extract of *Asparagus officinalis* stem, is a unique functional food that elicits anti-photoaging effects by suppressing pro-inflammatory signaling in hydrogen peroxide- and ultraviolet B-exposed skin fibroblasts. To elucidate its potential in preventing excessive inflammation in COVID-19, we examined the effects of ETAS®50 on pro-inflammatory responses in S1-stimulated murine peritoneal exudate macrophages. Co-treatment of the cells with ETAS®50 significantly attenuated S1-induced secretion of interleukin (IL)-6 in a concentration-dependent manner without reducing cell viability. ETAS®50 also markedly suppressed the S1-induced transcription of IL-6 and IL-1β. However, among the TLR4 signaling proteins, ETAS®50 did not affect the degradation of inhibitor κBα, nuclear translocation of nuclear factor-κB p65 subunit, and phosphorylation of c-Jun N-terminal kinase p54 subunit after S1 exposure. In contrast, ETAS®50 significantly suppressed S1-induced phosphorylation of p44/42 mitogen-activated protein kinase (MAPK) and Akt. Attenuation of S1-induced transcription of IL-6 and IL-1β by the MAPK kinase inhibitor U0126 was greater than that by the Akt inhibitor perifosine, and the effects were potentiated by simultaneous treatment with both inhibitors. These results suggest that ETAS®50 attenuates S1-induced IL-6 and IL-1β production by suppressing p44/42 MAPK and Akt signaling in macrophages. Therefore, ETAS®50 may be beneficial in regulating excessive inflammation in patients with COVID-19.

## 1. Introduction

Coronavirus disease 2019 (COVID-19) is an infectious disease caused by a novel type of coronavirus referred to as severe acute respiratory syndrome coronavirus 2 (SARS-CoV-2). Although 81% of infected patients develop either mild or uncomplicated illness, 14% develop pneumonia that requires hospitalization and oxygen inhalation, and 5% of patients become critically ill with respiratory failure, systemic shock, or multiple organ failure [1, 2]. In particular, people with a state of low-grade systemic chronic inflammation, such as advanced aging [3, 4], obesity [5–7], and type 2 diabetes [8–10], have a higher risk of death from COVID-19. Indeed, growing evidence suggests that excessive host inflammatory responses are associated with disease severity and mortality in patients [11, 12].

Severe cases demonstrated markedly high levels of tumor necrosis factor-α (TNF-α), interleukin (IL)-6, IL-10, and the soluble form of IL-2 receptor in circulation [13], exhibiting features similar to those of cytokine storm syndromes, such as macrophage activation syndrome [12]. Macrophages produce pro-inflammatory cytokines after detecting a broad range of pathogen-associated molecular patterns using pattern recognition receptors, such as toll-like receptors (TLRs). We recently reported that the SARS-CoV-2 spike protein S1 subunit (S1) strongly induces IL-6 and IL-1β production in murine and human macrophages by activating TLR4 signaling, similar to the action of pyrogenic lipopolysaccharide (LPS) [14]. Of note, recent clinical studies suggest that the IL-6 receptor antagonists tocilizumab and sarilumab and the IL-1 receptor antagonist anakinra improve the survival of patients with COVID-19 [15, 16]. Since manifestation of inflammatory disorders are influenced by an individual’s lifestyle, it is important to explore functional foods as prophylactic approach that can suppress excessive and undesired pro-inflammatory responses in macrophages.

ETAS®50 is a standardized extract of *Asparagus officinalis* stem, produced by Amino Up Co., Ltd. (Sapporo, Japan). Initially, it was discovered to be a novel and unique functional food that attenuates sleep deprivation-induced stress responses and promotes sleep in mice and humans [17, 18]. A subsequent study also reported that ETAS®50 intake reduced the feelings of dysphoria and fatigue, ameliorated the quality of sleep, enhanced stress-load performance, and increased salivary secretory immunoglobulin A levels in healthy adults [19]. Interestingly, a recent report showed that ETAS®50 supplementation restores cognitive functions and prevents neuronal apoptosis in the hippocampus of transgenic mice overexpressing amyloid precursor protein, which mimics Alzheimer’s disease [20]. The *in vivo* protective actions of ETAS®50 are suggested to be mediated by its ability to induce the expression of heat-shock protein 70 (HSP70) [17, 18, 20, 21], and asfral [22] and asparaprolines [23] have been identified as compounds that can induce HSP70 expression at the cellular and individual levels.

Moreover, our group previously reported that ETAS®50 attenuates hydrogen peroxide-induced expression of matrix metalloproteinase 9 and pro-inflammatory mediators by suppressing phosphorylation of c-Jun N-terminal kinase (JNK) and nuclear translocation of nuclear factor-κB (NF-κB) p65 subunit, respectively, in murine skin L929 fibroblasts [24, 25]. ETAS®50 also attenuated ultraviolet B-induced expression of IL-6 and IL-1β by suppressing the phosphorylation of Akt and nuclear translocation of p65, respectively, in normal human dermal fibroblasts [26, 27]. These previous findings suggest that ETAS®50 has the potential to abrogate pro-inflammatory responses by inhibiting TLR4 signaling in S1-stimulated macrophages. To elucidate its potential in preventing excessive inflammation in COVID-19, we examined the effects of ETAS®50 on pro-inflammatory responses in S1-stimulated murine primary macrophages.

## 2. Materials and Methods

### 2.1. Preparation of ETAS®50

ETAS®50, a standardized extract of *Asparagus officinalis* stem, was prepared from unused parts of asparagus in the factory of Amino Up Co., Ltd. (Sapporo, Japan) [17, 18, 22]. The asparagus is grown in Hokkaido, Japan and is commonly in the marketplace. In brief, *Asparagus officinalis* L. stems were collected and dried, and then extracted with hot water at 100 °C for 45 min. The extract was cooled to 50 °C and treated with cellulase and hemicellulase for 18 h to avoid clogging during production. After inactivation (100 °C, 20 min) of the enzymes, the extract was separated by centrifugation, concentrated *in vacuo*, and mixed with dextrin as a filler. The mixture was sterilized at 121 °C for 45 min and then spray-dried to produce ETAS®50 powder, consisting of 50% solid content of asparagus extract and 50% dextrin. Component analysis showed that the ETAS®50 powder comprised 78.5% carbohydrates, 12.3% proteins, 5.0% ashes, 0.7% lipids, and 3.5% moisture. The manufacturing process was conducted in accordance with good manufacturing practice standards for dietary supplements and ISO9001:2015 and ISO22000:2018 criteria.

### 2.2. Animal care and use

Adult (8–12-week-old) male C57BL/6J mice (Sankyo Labo Service, Tokyo, Japan) were housed at a temperature of 23–25 °C and humidity of 50%–60% with a fixed light/dark cycle (light, 7:00–19:00; dark, 19:00–7:00). Food and water were provided ad libitum. This study was approved by the Experimental Animal Ethics Committee in Kyorin University (No. 245, 2021). All experiments described below were carried out following the Guiding Principles for the Care and Use of Animals approved by the Council of the Physiological Society of Japan, in accordance with the Declaration of Helsinki, 1964.

### 2.3. Preparation and culture of peritoneal exudate macrophages

Two milliliters of sterilized 4.05% thioglycollate medium Brewer modified (Becton, Dickinson and Company, Franklin Lakes, NJ, USA) was administered intraperitoneally into the mice, and the mice were housed for four days [14, 28]. After the mice were euthanized by cervical dislocation, peritoneal exudate cells were harvested by sterile lavage of the peritoneal cavity with ice -cold Dulbecco’s modified Eagle’s medium (DMEM; Nacalai Tesque, Kyoto, Japan). The cells were washed once with ice-cold DMEM, resuspended in DMEM supplemented with 1 0% heat-inactivated fetal bovine serum (BioWest, Nuaillé, France), 100 units/ml penicillin (Nacalai Tesque), and 100 μg/ml streptomycin (Nacalai Tesque), and then cultured at 37 °C in a humidified incubator containing 5% CO_2_ for 1 h. After the nonadherent cells were removed, peritoneal exudate macrophages were used in the experiments.

### 2.4. Agents and treatment

Different concentrations of ETAS®50 or dextrin (vehicle control) (0.25, 0.5, 1, or 2 mg/mL) were prepared by directly dissolving each agent in the complete medium, and then the supplemented medium was filter sterilized using a 0.22 μm membrane [21, 24–27]. To assess its anti-inflammatory effects, the cells were co-treated with the indicated concentrations of ETAS®50 or dextrin and 100 ng/mL of SARS-CoV-2 spike recombinant protein S1 subunit (Arigo Biolaboratories, Hsinchu City, Taiwan) for 1 to 24 h. The cells were treated with 20 μM nigericin (Sigma-Aldrich, St. Louis, MO, USA) for the last 1 h of S1 stimulation to promote IL-1β processing and secretion. Cells were co-treated with 5 μM U0126 (Cell Signaling Technology, Danvers, MA, USA) or 20 μM perifosine (Cell Signaling Technology) to block S1-induced phosphorylation of p44/42 MAPK or Akt, respectively. The final concentrations of vehicles used to dissolve these agents were equivalent to the culture medium used among the experimental groups.

### 2.5. Enzyme-linked immunosorbent assay (ELISA)

Cell culture supernatants were collected after centrifugation at 300 × *g* for 20 min. Concentrations of IL-6 and IL-1β were measured using the Quantikine Mouse IL-6 ELISA Kit (R&D Systems, Minneapolis, MN, USA) and Quantikine Mouse IL-1β ELISA Kit (R&D Systems), as described previously [14, 28]. Since the detection limits of IL-6 and IL-1β for ELISA are 7.8–500 pg/mL and 12.5–800 pg/mL, respectively, the supernatants were diluted 10 times before measurements.

### 2.6. Cell viability assay

Cell viability was measured using the Cell Counting Kit-8 (FUJIFILM Wako Pure Chemical, Osaka, Japan) [29]. After cell culture supernatants were collected, fresh complete medium supplemented with 10% Cell Counting Kit-8 reagent was added to each well, and incubation was continued for 1 h. After the reaction was stopped by adding 1% sodium dodecyl sulfate, the absorbance of medium in each well was analyzed using a multi-mode microplate reader FilterMax F5 (Molecular Devices, San Jose, CA, USA) at a wavelength of 450 nm. Cell viability was calculated with the viability of vehicle-treated control cells as 100%.

### 2.7. Reverse transcription and real-time polymerase chain reaction (PCR)

Total cellular RNA was extracted using RNAiso Plus reagent (TaKaRa Bio, Shiga, Japan). One microgram of total cellular RNA was converted to single-stranded cDNA using the PrimeScript 1st Strand cDNA Synthesis Kit (Takara Bio). The cDNA (1 μl) was amplified using Premix Ex Taq (Probe qPCR) (TaKaRa Bio) in a 7500 Real Time PCR System (Thermo Fisher Scientific, Waltham, MA, USA). The PCR conditions were as follows: 50 °C for 2 min and 95 °C for 15 s, followed by 40 cycles of 95 °C for 15 s and 60 °C for 1 min. The fluorescent probes and primers used are listed in Table 1. The mRNA expression levels of target genes were calculated as the ratio of their values to that of 18S rRNA as an internal control.

**Table 1.**
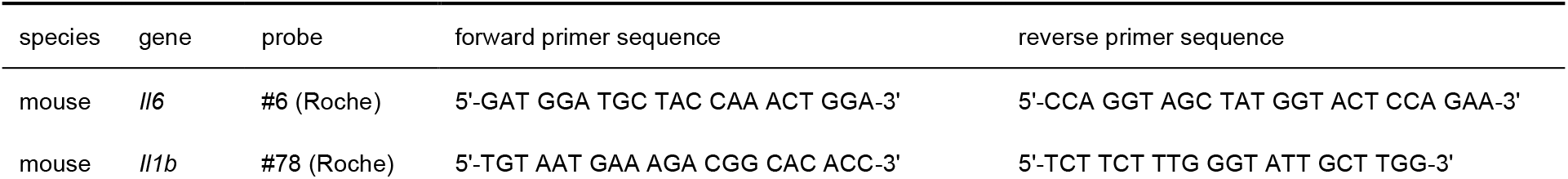
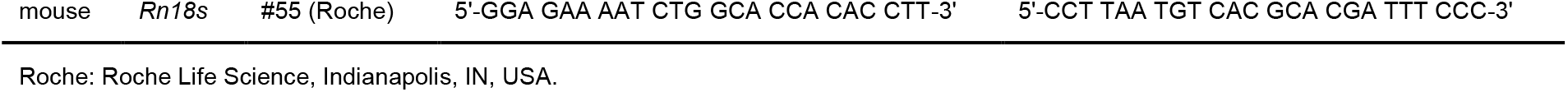
Fluorescent probes and primers used in this study

### 2.8. Preparation of nuclear extracts

Nuclear proteins were prepared as previously described [14, 25, 26]. The cells were extracted in lysis buffer containing 10 mM HEPES–KOH (pH 7.8), 10 mM KCl, 2 mM MgCl_2_, 0.1 mM ethylenediaminetetraacetic acid (EDTA), and 0.1% Nonidet P-40 supplemented with protease and phosphatase inhibitor cocktails (Nacalai Tesque). After low-speed centrifugation (200 × *g*) at 4 °C for 5 min, sediments containing the nuclei were resuspended in wash buffer containing 250 mM sucrose, 10 mM HEPES–KOH (pH 7.8), 10 mM KCl, 2 mM MgCl_2_, and 0.1 mM EDTA. The suspension was centrifuged at low-speed (200 ×*g*) at 4 °C for 5 min, then the sediments were resuspended in nuclear extraction buffer containing 50 mM HEPES–KOH (pH 7.8), 420 mM KCl, 5 mM MgCl_2_, 0.1 mM EDTA, and 20% glycerol, and rotated at 4 °C for 30 min. After high-speed centrifugation (13,000 × *g*) at 4 °C for 15 min, the supernatants were used as source of nuclear proteins. The concentration of nuclear proteins was determined using the Protein Assay BCA kit (Nacalai Tesque).

### 2.9. Western blotting

Whole or nuclear proteins (10 μg) were separated through electrophoresis on a sodium dodecyl sulfate-polyacrylamide gel and then transferred onto an Immobilon-P membrane (Merck Millipore, Burlington, MA, USA). After blocking with 5% bovine serum albumin, each membrane was probed with primary antibodies for 1 h (Table 2). Secondary antibodies conjugated with horseradish peroxidase (Jackson ImmunoResearch Laboratories, West Grove, PA, USA) were applied at a 1:20,000 dilution for 30 min. The membrane was incubated with Western BLoT Chemiluminescence HRP Substrate (TaKaRa Bio), and membrane image was captured using a chemiluminescent imaging system LuminoGraph I (ATTO, Tokyo, Japan). The density of each protein band was quantified using ImageJ software (National Institutes of Health, Bethesda, MD, USA). Glyceraldehyde-3-phosphate dehydrogenase (GAPDH) and Yin Yang 1 (YY1) were used as loading controls for whole and nuclear proteins, respectively. The phosphorylation levels of the target proteins were calculated as the ratio of their values to those of the corresponding total protein as a loading control.

**Table 2.** Primary antibodies used in this study

| protein | M.W. | Cat. No. | manufacturer | Dilution | protein | M.W. | Cat. No. | manufacturer | dilution |
| --- | --- | --- | --- | --- | --- | --- | --- | --- | --- |
| IκBα | 39 kDa | 4814S | CST | 1/1,000 | GAPDH | 37 kDa | 5174S | CST | 1/2,000 |
| p65 | 65 kDa | 8242S | CST | 1/1,000 | YY1 | 68 kDa | ab109237 | Abcam | 1/2,000 |
| p-JNK | 54/46 kDa | 4668S | CST | 1/1,000 | JNK | 54/46 kDa | 9258S | CST | 1/1,000 |
| p-p44/42 | 44/42 kDa | 4370P | CST | 1/1,000 | p44/42 | 44/42 kDa | 4695P | CST | 1/1,000 |
| p-Akt | 60 kDa | 4060S | CST | 1/1,000 | Akt | 60 kDa | 4691S | CST | 1/1,000 |

CST: Cell Signaling Technology, Danvers, MA, USA.

### 2.10. Fluorescence immunomicroscopy

The cells were fixed with 4% paraformaldehyde for 15 min and then permeabilized with methanol at -20 °C for 10 min. After blocking with 1% bovine serum albumin, the primary antibody against p65 (8242S; Cell Signaling Technology) was applied at a 1:400 dilution for 1 h. Then, the secondary antibody conjugated with Alexa Fluor 568 (Abcam, Cambridge, UK) was applied at a 1:1,000 dilution with 3 μM Nuclear Green DCS1 (Abcam) for 30 min. After mounting on glass slides, the subcellular localization of p65 and the nucleus was visualized with FL2 (orange-red) and FL1 (green) detectors, respectively, using Nikon BioStation IM (NIKON, Tokyo, Japan) [14, 26].

### 2.11. Statistical analysis

Experimental data are presented as mean ± standard error of the mean (SEM). Comparisons among at least three groups were tested using one-way analysis of variance (ANOVA), and post hoc comparisons to determine significant differences between two groups were performed using the Bonferroni test. Differences were considered statistically significant at *p* < 0.05.

## 3. Results

### 3.1. ETAS®50 attenuated S1-induced IL-6 secretion in a concentration-dependent manner without reducing the cell viability of macrophages

To elucidate whether ETAS®50 has anti-inflammatory effects on S1-stimulated macrophages, we first examined the concentration-dependent effects of ETAS®50 treatment on S1-induced secretion of IL-6 in murine peritoneal exudate macrophages. When the cells were simultaneously treated with different concentrations (0, 0.25, 0.5, 1, or 2 mg/mL) of ETAS®50 and 100 ng/mL of S1 for 6 h, ETAS®50 significantly attenuated S1-induced secretion of IL-6 in a concentration-dependent manner (Figure 1(a)). To rule out that the attenuation of the S1-induced IL-6 section by ETAS®50 is due to a reduction in the number of living cells, we conducted a cell viability assay under the same experimental conditions. The cells stimulated with S1 alone showed significant increase in cell viability, which was not influenced by ETAS®50 treatment (Figure 1(b)).

**FIGURE 1:**
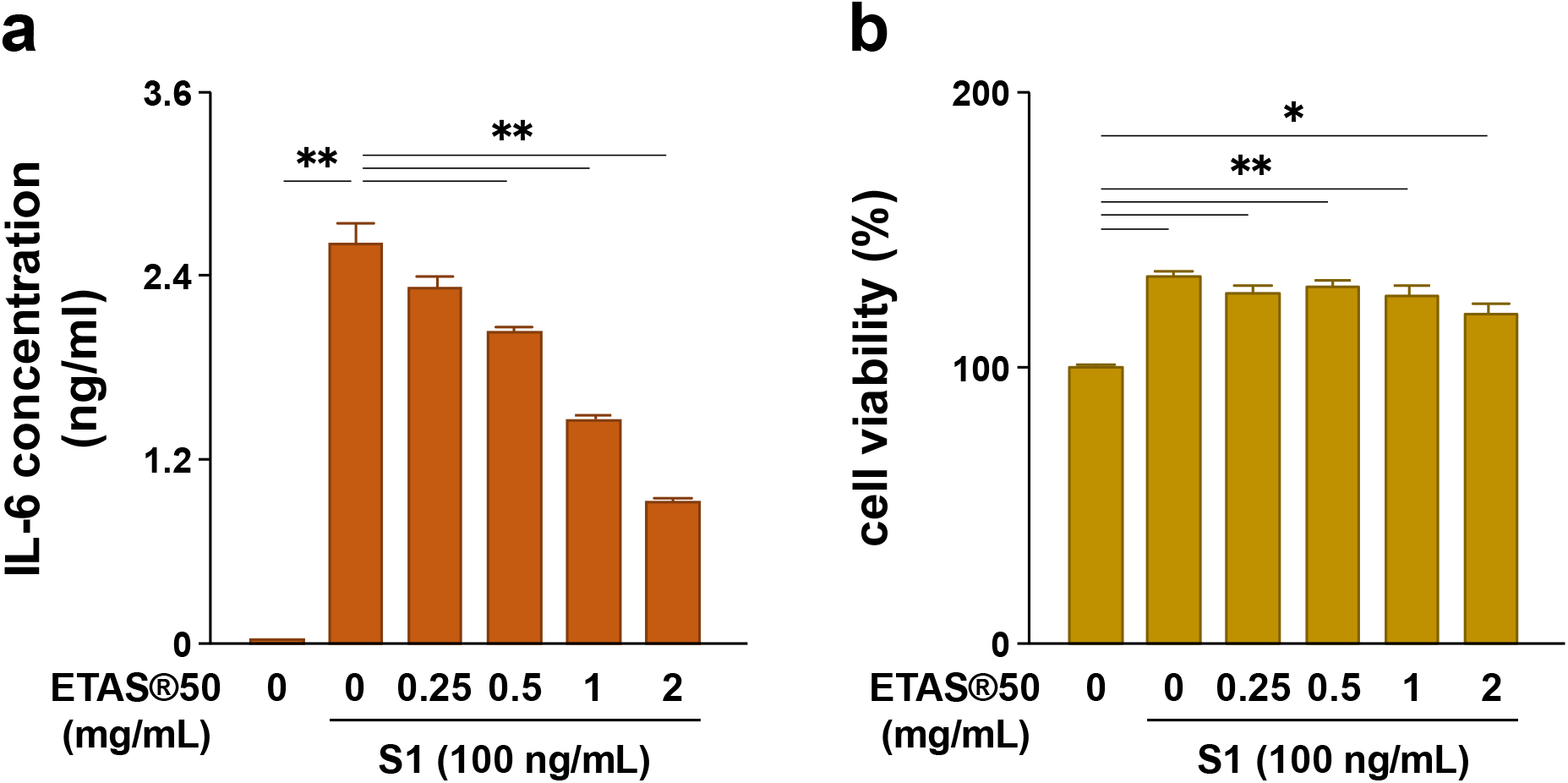
Effect of ETAS®50 on S1-induced secretion of IL-6 and viability of murine peritoneal exudate macrophages. The cells were co-treated with 0, 0.25, 0.5, 1, or 2 mg/mL of ETAS®50 and 100 ng/mL of S1 for 6 h. (a) IL-6 concentrations in culture supernatants were analyzed using ELISA. (b) Cell viability was analyzed using the Cell Counting Kit-8. Mean ± SEM (*n* = 3). \**p* < 0.05, \*\**p* < 0.01, using one-way ANOVA and Bonferroni test.

### 3.2. ETAS®50 repressed S1-induced IL-6 and IL-1β transcription in macrophages

To confirm that the ETAS®50 treatment attenuates S1-induced secretion of IL-6 by transcriptional repression, we next analyzed the effects of ETAS®50 treatment on S1-induced transcription of IL-6 and IL-1β in murine peritoneal exudate macrophages. When the cells were simultaneously treated with 2 mg/mL of ETAS®50 and 100 ng/mL of S1 for 6 h, ETAS®50 dramatically repressed S1-induced transcription of IL-6 and IL-1β (Figure 2(a)). Moreover, extending the treatment time from 6 h to 24 h potentiated the attenuating effect of ETAS®50 on S1-induced secretion of IL-6 (Figure 1(a) and 2(b): left). Moreover, ETAS®50 strongly attenuated nigericin-induced secretion of IL-1β after priming the cells with S1 stimulation for 24 h (Figure 2(b): right).

**FIGURE 2:**
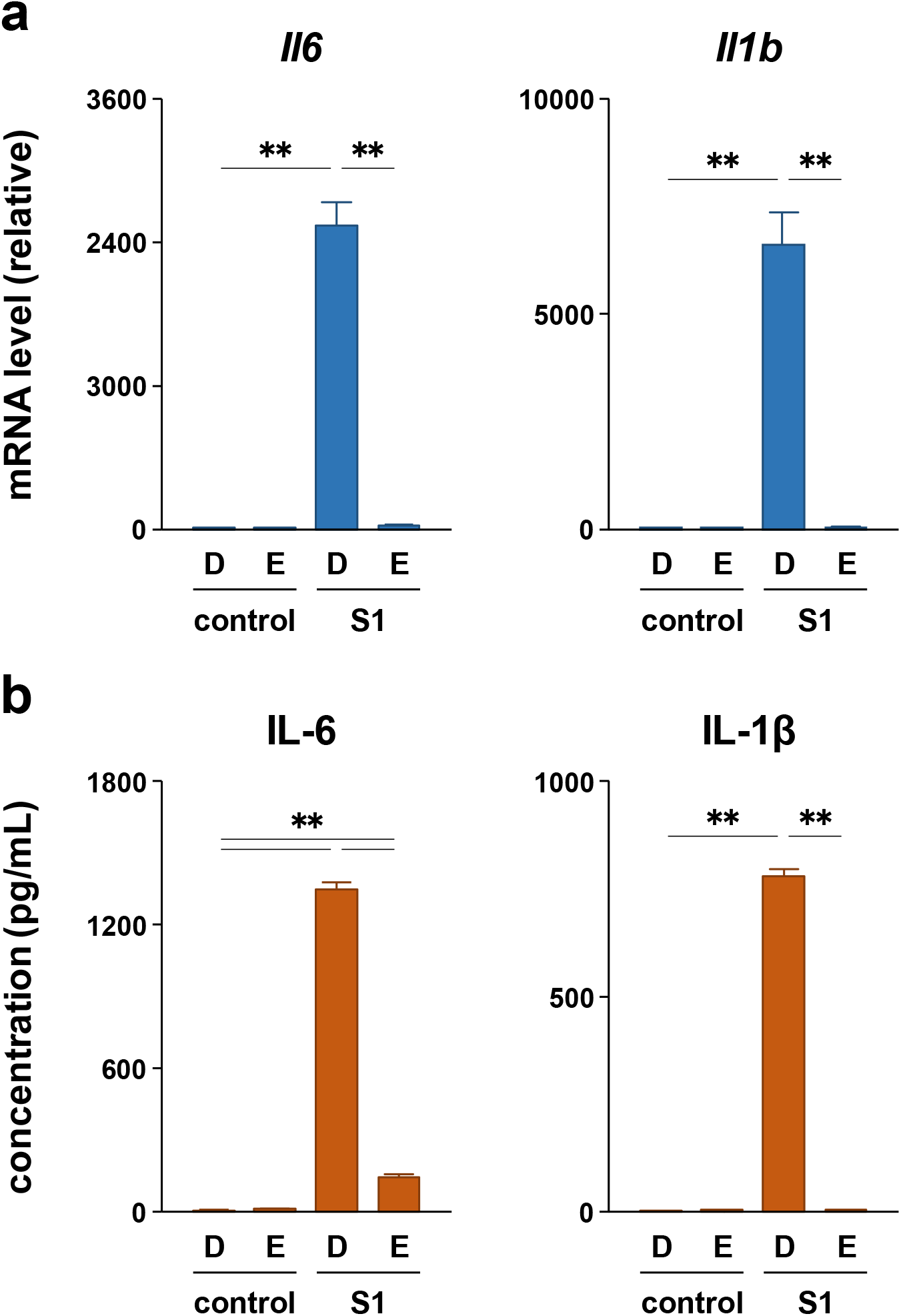
Effect of ETAS®50 on S1-induced transcription of IL-6 and IL-1β in murine peritoneal exudate macrophages. The cells were co-treated with 2 mg/mL of ETAS®50 (E) or dextrin (D; vehicle control) and 100 ng/mL of S1 for 6 h (a) and 24 h (b). (a) IL-6 (left) and IL-1β (right) mRNA levels were analyzed using real-time PCR. (b) IL-6 (left) and IL-1β (right) concentrations in culture supernatants were analyzed using ELISA. The cells were treated with 20 μM nigericin for the last 1 h of S1 stimulation to promote IL-1β processing and secretion (b: right). Mean ± SEM (*n* = 3). \*\**p* < 0.01, using one-way ANOVA and Bonferroni test.

### 3.3. ETAS®50 suppressed S1-induced p44/42 MAPK and Akt phosphorylation without affecting NF-κB nuclear translocation and JNK phosphorylation in macrophages

We recently reported that S1-induced transcription of pro-inflammatory cytokines is regulated by NF-κB and JNK signaling [14]. Therefore, we analyzed whether ETAS®50 suppresses S1-induced activation of NF-κB and JNK signaling in murine peritoneal exudate macrophages. Western blotting revealed that co-treatment of the cells with ETAS®50 (2 mg/mL) for 1 h did not affect S1 (100 ng/mL)-induced degradation of inhibitor κBα (IκBα), nuclear accumulation of NF-κB p65 subunit, and phosphorylation of JNK p54 subunit (Figure 3(a)). Immunofluorescence data also supported the result that ETAS®50 did not inhibit the S1-induced nuclear translocation of p65 (Figure 3(b)). However, instead of inhibiting the activation of NF-κB and JNK signaling, ETAS®50 significantly suppressed S1-induced phosphorylation of p44/42 MAPK and Akt after 6 h of co-treatment without interfering with basal phosphorylation levels (Figure 4).

**FIGURE 3:**
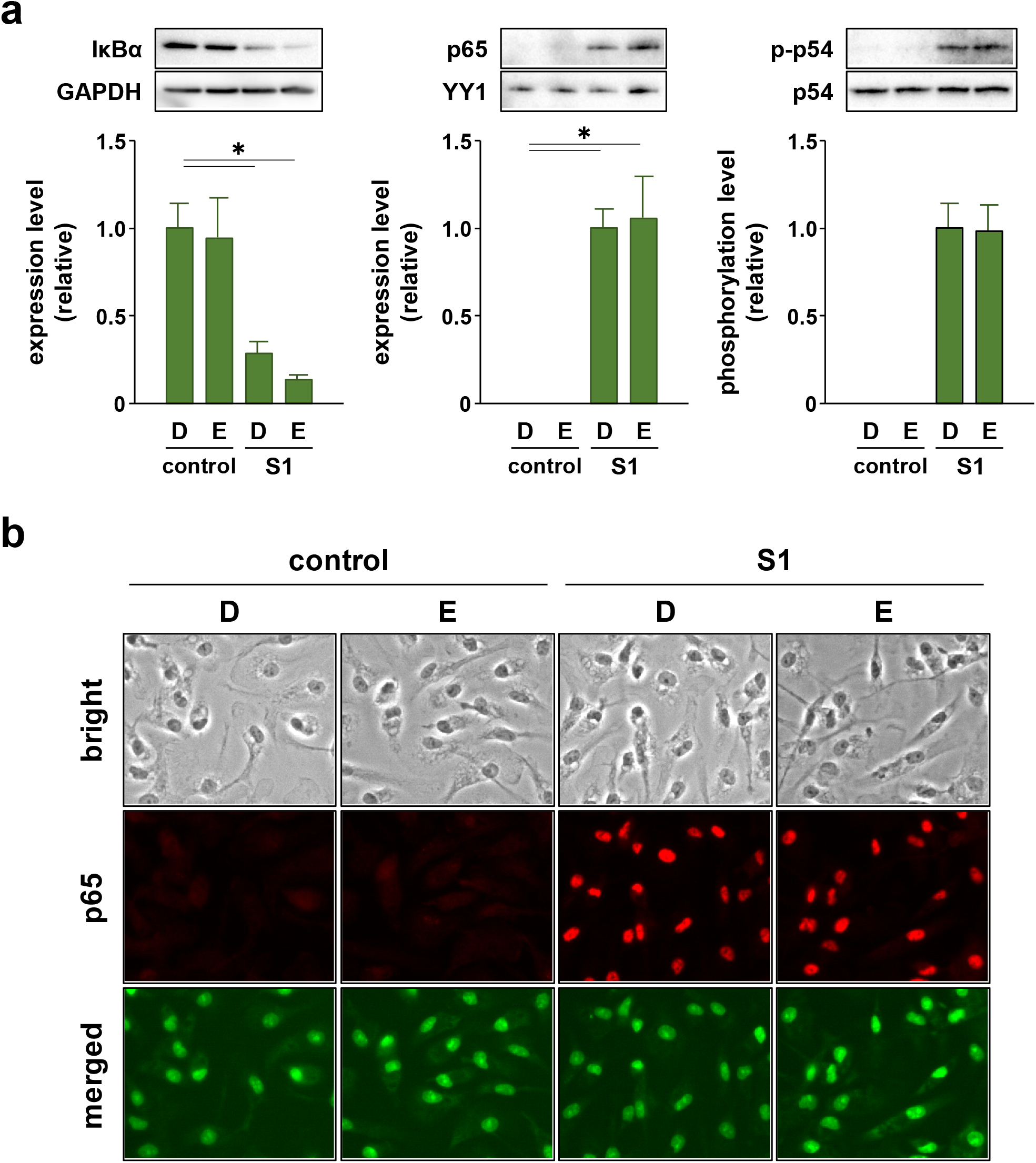
Effect of ETAS®50 on S1-induced activation of the NF-κB and JNK signaling in murine peritoneal exudate macrophages. The cells were co-treated with 2 mg/mL of ETAS®50 (E) or dextrin (D; vehicle control) and 100 ng/mL of S1 for 1 h. (a) Total amount of IκBα (left), nuclear amount of NF-κB p65 subunit (middle), and phosphorylation level of JNK p54 subunit (right) were analyzed using western blotting. Mean ± SEM (*n* = 3). \**p* < 0.05, using one-way ANOVA and Bonferroni test. (b) Subcellular localization of p65 was visualized using fluorescence immunomicroscopy.

**FIGURE 4:**
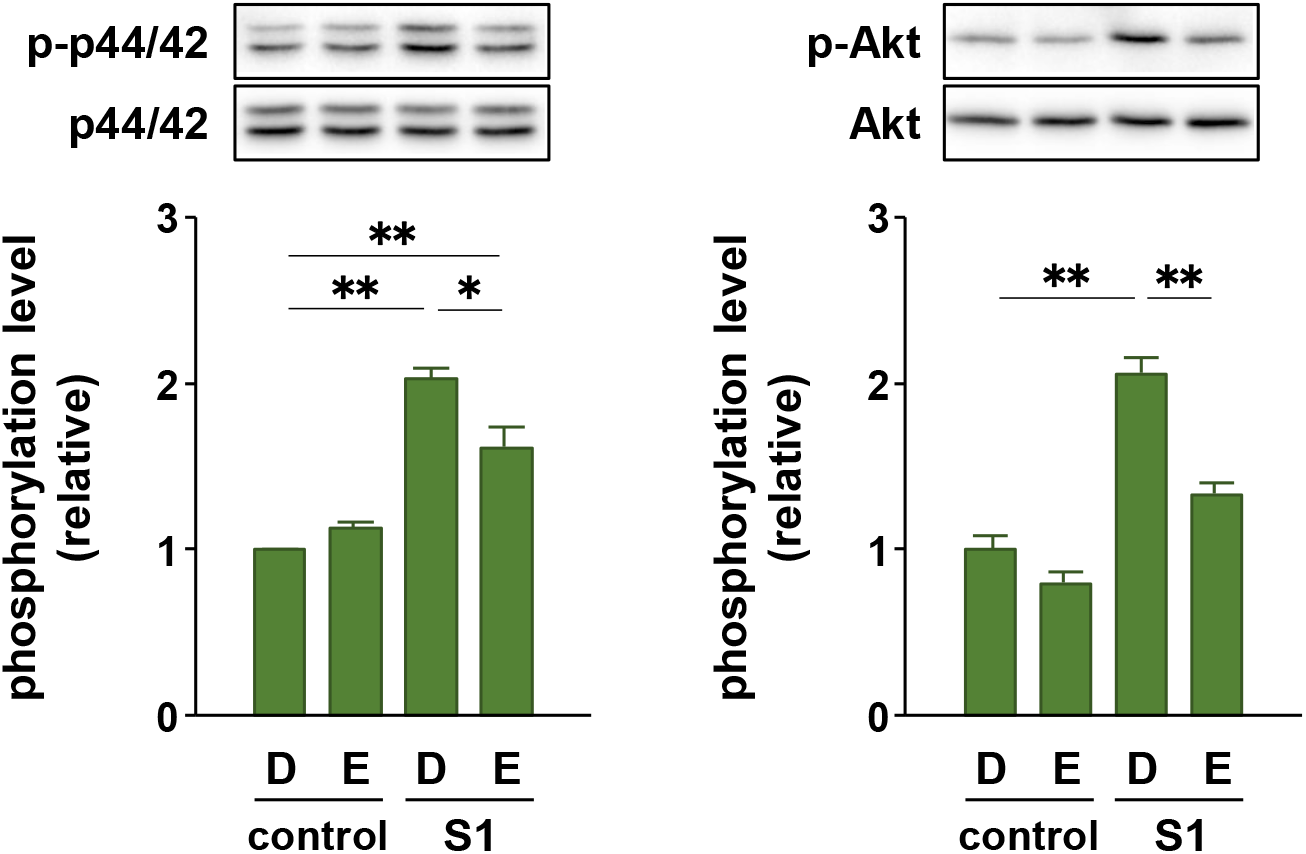
Effect of ETAS®50 on S1-induced phosphorylation of p44/42 MAPK and Akt in murine peritoneal exudate macrophages. The cells were co-treated with 2 mg/mL of ETAS®50 (E) or dextrin (D; vehicle control) and 100 ng/mL of S1 for 6 h. Phosphorylation levels of p44/42 MAPK (left) and Akt (right) were analyzed using western blotting. Mean ± SEM (*n* = 3). \**p* < 0.05, \*\**p* < 0.01, using one-way ANOVA and Bonferroni test.

### 3.4. The p44/42 MAPK and Akt signaling are additively involved in S1-induced IL-6 and IL-1β transcription in macrophages

To clarify whether p44/42 MAPK and Akt signaling mediate S1-induced pro-inflammatory responses, we examined the effects of the MAPK kinase inhibitor U0126 and Akt inhibitor perifosine on S1-induced transcription of IL-6 and IL-1β in murine peritoneal exudate macrophages. Co-treatment of the cells with 5 μM U0126 abolished S1-induced phosphorylation of p44/42 MAPK after 6 h (Figure 5(a)). At the same time, S1-induced transcription of IL-6 was significantly decreased by half, and that of IL-1β was dramatically suppressed (Figure 5(b)). On the other hand, co-treatment of the cells with 20 μM perifosine significantly mitigated S1-induced phosphorylation of Akt after 6 h (Figure 6(a)), which was accompanied by significant repression of S1-induced transcription of IL-1β, but not of IL-6 (Figure 6(b)). When the cells were simultaneously treated with U0126 and perifosine to mimic the condition treated with ETAS®50, transcriptional repression of IL-6 and IL-1β was augmented compared to treatment with each inhibitor alone (Figure 7).

**FIGURE 5:**
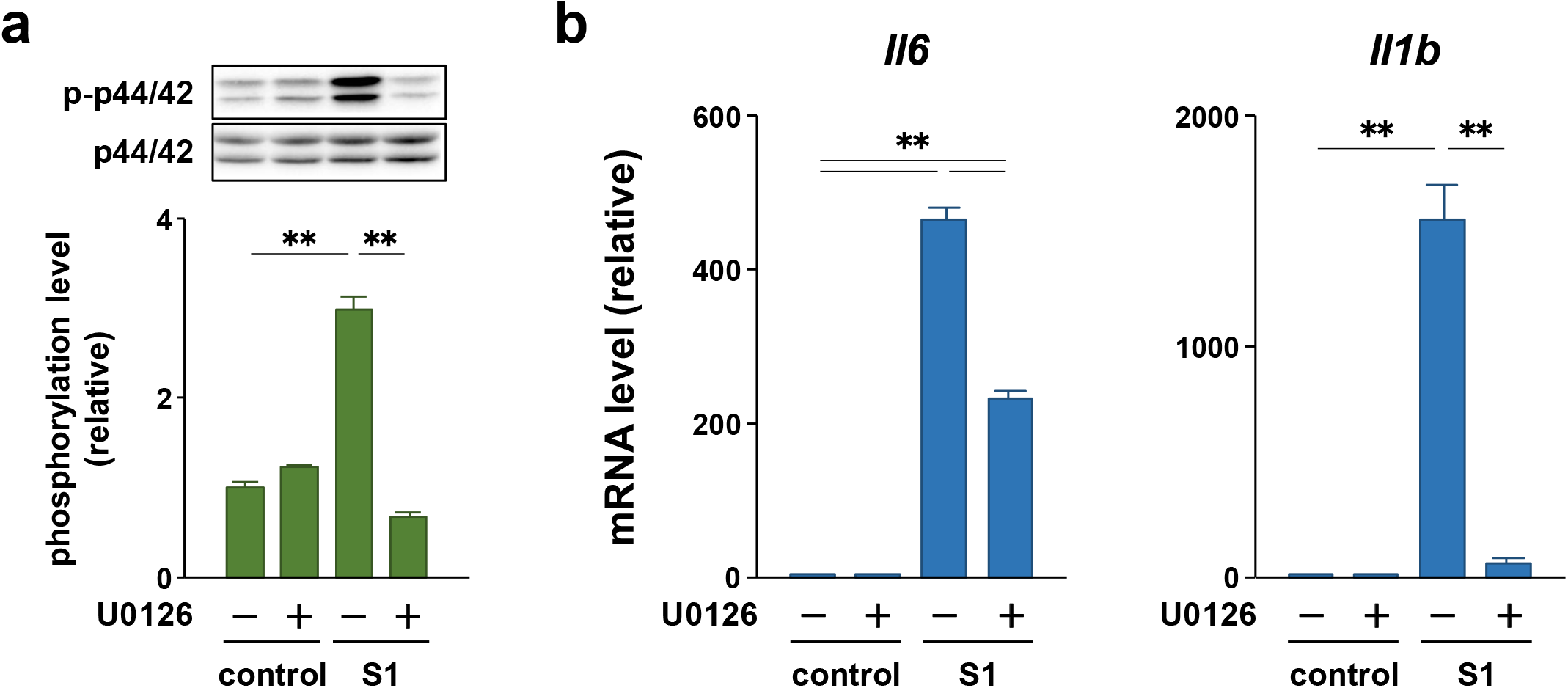
Effect of the MAPK inhibitor U0126 on S1-induced transcription of IL-6 and IL-1β in murine peritoneal exudate macrophages. The cells were co-treated with 5 μM U0126 or dimethyl sulfoxide alone (vehicle control) and 100 ng/mL of S1 for 6 h. (a) Phosphorylation level of p44/42 MAPK was analyzed using western blotting. (b) IL-6 (left) and IL-1β (right) mRNA levels were analyzed using real-time PCR. Mean ± SEM (*n* = 3). \*\**p* < 0.01, using one-way ANOVA and Bonferroni test.

**FIGURE 6:**
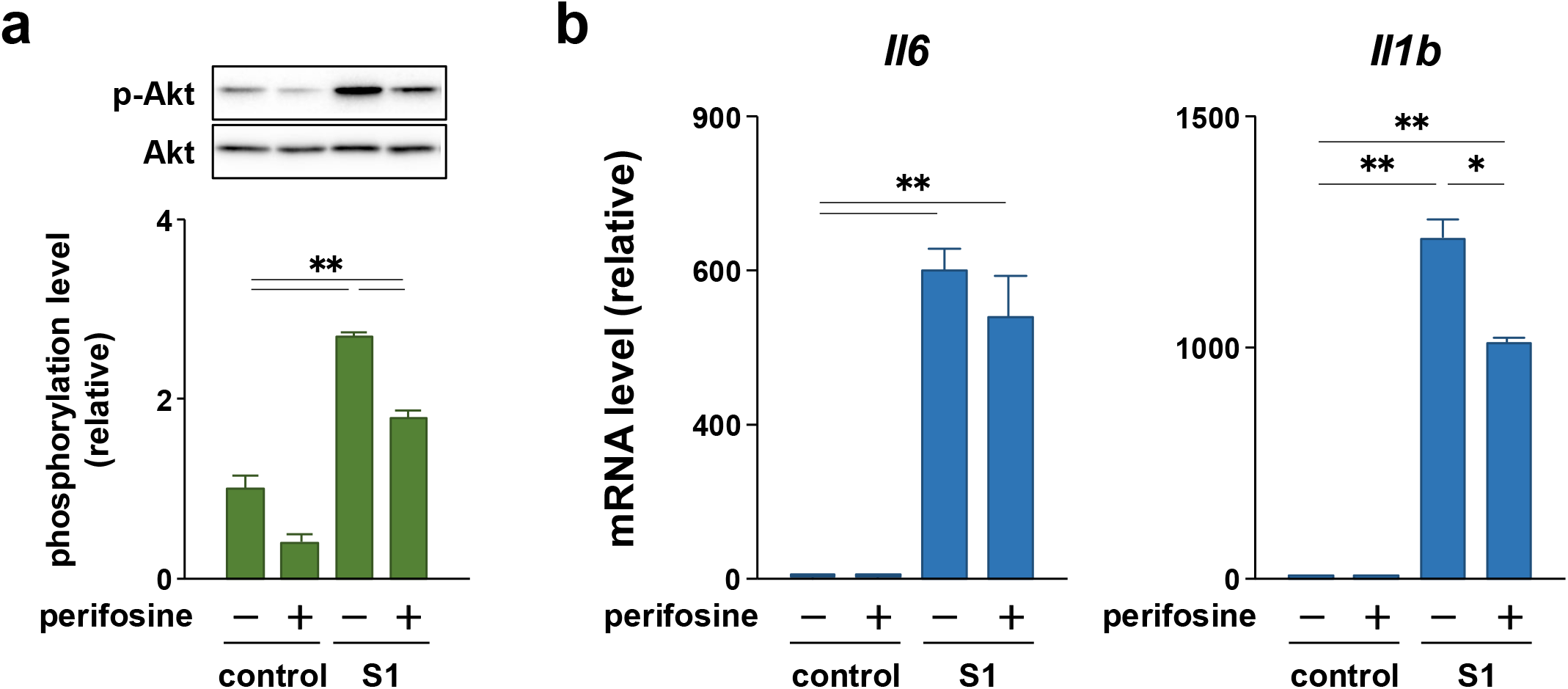
Effect of the Akt inhibitor perifosine on S1-induced transcription of IL-6 and IL-1β in murine peritoneal exudate macrophages. The cells were co-treated with 20 μM perifosine or sterile water alone (vehicle control) and 100 ng/mL of S1 for 6 h. (a) Phosphorylation level of Akt was analyzed using western blotting. (b) IL-6 (left) and IL-1β (right) mRNA levels were analyzed using real-time PCR. Mean ± SEM (*n* = 3). \**p* < 0.05, \*\**p* < 0.01, using one-way ANOVA and Bonferroni test.

**FIGURE 7:**
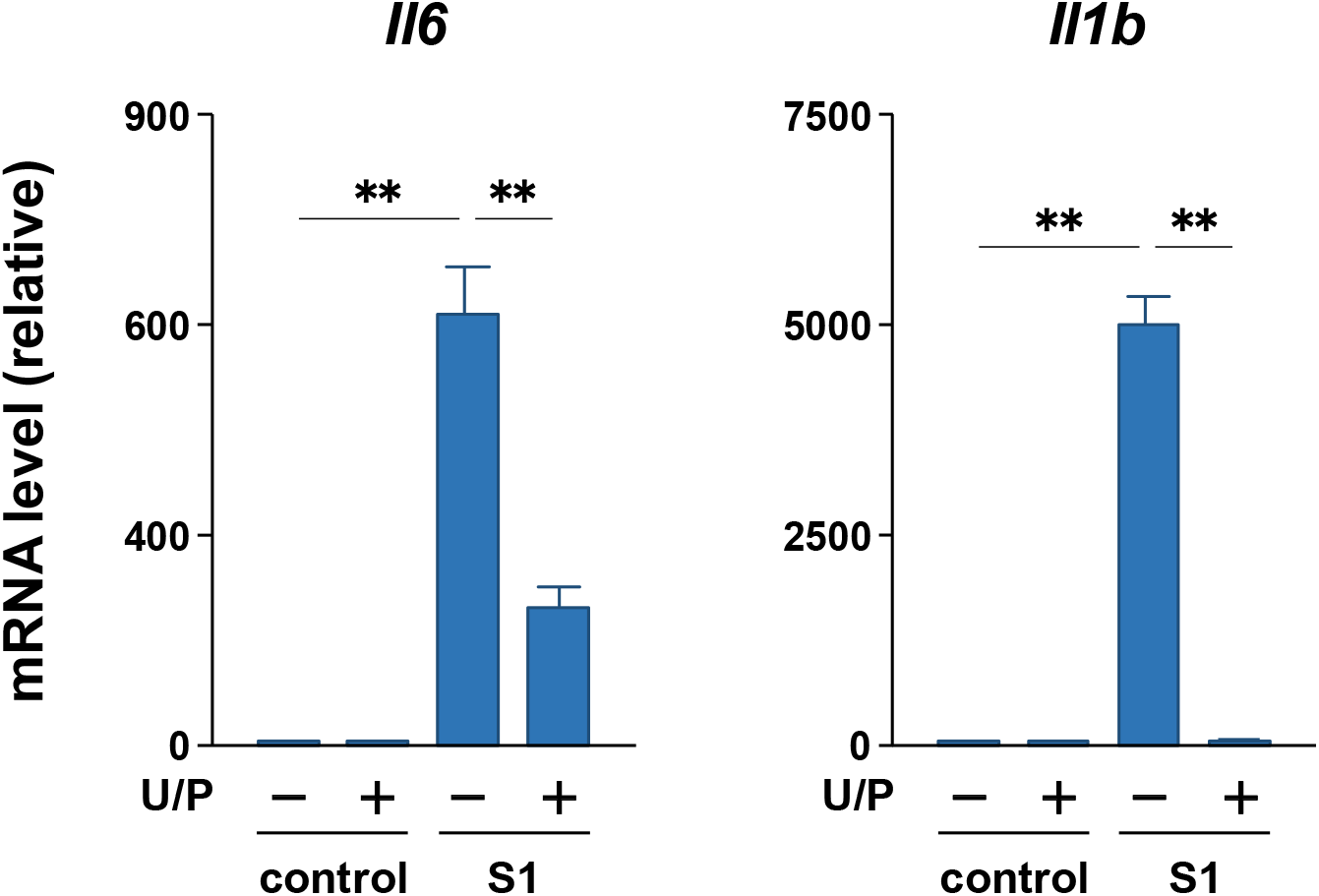
Effect of simultaneous treatment with U0126 and perifosine on S1-induced transcription of IL-6 and IL-1β in murine peritoneal exudate macrophages. The cells were co-treated with 5 μM U0126 (U), 20 μM perifosine (P), and 100 ng/mL of S1 for 6 h. The concentration of vehicles is same across all samples. IL-6 (left) and IL-1β (right) mRNA levels were analyzed using real-time PCR. Mean ± SEM (*n* = 3). \*\**p* < 0.01, using one-way ANOVA and Bonferroni test.

## 4. Discussion

The pro-inflammatory cytokine IL-6 has been suggested to be involved in the aggravation of patients with COVID-19 [15, 16]. In the initial experiment, we found that co-treatment with ETAS®50 attenuated S1-induced secretion of IL-6 in murine peritoneal exudate macrophages in a concentration-dependent manner after 6 h of culture. In addition, this result is not due to a reduction in the number of living cells, since ETAS®50 did not influence cell viability. The viability of S1-treated cells was significantly higher than that in vehicle-treated control cells. However, primary cultured macrophages were terminally differentiated and therefore cannot undergo cell proliferation. To analyze the number of viable cells, we used a water-soluble tetrazolium salt WST-1, which was reduced to formazan dye by the activity of mitochondrial dehydrogenases in metabolically active cells. A previous study has demonstrated that the TLR4 agonist LPS facilitates mitochondrial oxidation by succinate dehydrogenase in bone marrow-derived macrophages [30]. LPS also promoted reduction of thiazolyl blue tetrazolium bromide MTT into formazan dye in RAW264.7 macrophages [31]. Therefore, our results suggest that S1 activates the enzymes in macrophages as TLR4 agonist in a manner similar to the action of LPS, and that ETAS®50 can attenuate S1-induced production of IL-6 without impairing cell survival and metabolism.

The attenuation of S1-induced IL-6 secretion is due to its transcriptional repression, since ETAS®50 co-treatment dramatically repressed S1-induced transcription of IL-6 after 6 h of culture. Moreover, when the treatment time was extended from 6 h to 24 h, the attenuating effect of ETAS®50 on S1-induced IL-6 secretion was clearly potentiated. These results suggest that ETAS®50 has a weak or no suppressive effect on S1-induced signal transduction in the early phase, such as NF-κB and JNK signaling, which regulates the transcription of IL-6. We recently reported that S1 induces IκBα degradation, NF-κB p65 subunit nuclear translocation, and JNK phosphorylation by interacting with TLR4 in macrophages within 1 h of culture [14]. Indeed, in this study, co-treatment with ETAS®50 did not suppress the activation of NF-κB and JNK signaling by S1 exposure. In contrast, ETAS®50 treatment suppressed NF-κB p65 nuclear translocation and JNK phosphorylation in hydrogen peroxide- or ultraviolet B-exposed skin fibroblasts [24–26]. The discrepancy in the action mechanisms of ETAS®50 may be due to the difference in upstream signal transduction pathways between TLR4 and other stress-triggered signaling pathways.

It has been also suggested that suppressing the action of IL-1 may be a therapeutic target for critically-ill patients with COVID-19 [15]. In this study, ETAS®50 co-treatment dramatically repressed S1-induced transcription of IL-1β. The regulatory machinery for IL-1β processing and secretion substantially differs from that of most other pro-inflammatory cytokines, including IL-6. IL-1β is initially synthesized as a leaderless precursor that requires cleavage into its active form by inflammasome-activated caspase-1 [32]. The inflammasome is a multimeric protein complex consisting of the Nod-like receptor family pyrin domain containing 3 (NLRP3), apoptosis-associated speck-like protein containing a caspase recruitment domain, and pro-caspase-1, whose formation is facilitated by endogenous danger-associated molecular patterns (DAMPs), such as ATP, cholesterol crystals, urate crystals, and ceramide [33]. Growing evidence suggests that NLRP3 inflammasome activation in macrophages and lung epithelial cells is involved in the development of pneumonia in patients with COVID-19 [34–36]. In this study, ETAS®50 strongly attenuated the DAMP nigericin-induced secretion of IL-1β after priming the cells with S1 stimulation for 24 h. Therefore, ETAS®50 attenuated both S1-induced IL-6 and IL-1β production by repressing their transcription in macrophages.

In addition to NF-κB and JNK signaling, the activation of p44/42 MAPK and Akt signaling was observed to regulate the transcription of pro-inflammatory cytokines in macrophages. Indeed, it has been reported that a wide variety of herbal extracts and herb-derived natural compounds exert anti-inflammatory effects on LPS-stimulated macrophage cell lines, such as RAW264.7 and U937, by suppressing either p44/42 MAPK signaling [37– 40] or Akt signaling [41–43]. In this regard, we found that ETAS®50 co-treatment suppressed S1-induced phosphorylation of p44/42 MAPK and Akt after 6 h of culture. Moreover, S1-induced transcription of IL-6 and IL-1β was largely regulated by p44/42 MAPK signaling in murine peritoneal exudate macrophages, since the transcriptional induction was significantly repressed by co-treatment with the MAPK kinase inhibitor U0126. In contrast, Akt signaling had a minor contribution to the transcriptional induction of IL-6 and IL-1β by S1 exposure compared to p44/42 MAPK signaling, as the Akt inhibitor perifosine partially repressed IL-1β transcription but did not influence IL-6 transcription.

Simultaneous suppression of S1-induced phosphorylation of p44/42 MAPK and Akt augmented the repression of S1-induced IL-6 and IL-1β transcription, suggesting that ETAS®50 dramatically attenuated S1-induced IL-6 and IL-1β production by simultaneously suppressing both p44/42 MAPK and Akt signaling. However, the unknown actions of ETAS®50 should be considered, because IL-6 transcription could not be dramatically repressed even when both S1-induced p44/42 and Akt phosphorylation was largely suppressed by the inhibitors.

More importantly, ETAS®50 did not interfere with the basal phosphorylation of p44/42 MAPK and Akt in murine primary macrophages. p44/42 MAPK is also a signal transduction protein downstream of receptor tyrosine kinases and integrins that regulate cellular motility [44]. Akt signaling acts as a master regulator for maintaining cellular proliferation, survival, and metabolism downstream of growth factor receptors [45]. Therefore, it may be extremely important for preventing impairment of macrophage functions that ETAS®50 does not indiscriminately suppress p44/42 MAPK and Akt phosphorylation. These results are also consistent with the capability of ETAS®50 to exert the attenuating effect on S1-induced IL-6 and IL-1β production in macrophages without reducing the number of living cells. Acute and subacute ETAS®50 administration had no significant side effects on food consumption, body weight, mortality, urinalysis, hematology, biochemistry, necropsy, organ weight, and histopathology in rats [46], while it can exert beneficial effects such as anti-stress, promoting sleep, and improving cognitive impairment in mice and humans [17–20]. Thus, the ability of ETAS®50 to exert various beneficial effects without side effects *in vivo* may be related to its failure to suppress signal transduction, which is essential for maintaining cellular functions.

## 4. Conclusion

Excessive host inflammatory responses are associated with disease severity and mortality in patients with COVID-19. In particular, suppressing IL-6 and IL-1 production and blocking their receptors and actions are currently attracting attention as therapeutic targets for severe COVID-19. Growing evidence also suggests that the SARS-CoV-2 spike protein contributes to the development of macrophage activation syndrome by activating TLR4 signaling. In this study, ETAS®50 attenuated S1-induced IL-6 and IL-1β production by suppressing p44/42 MAPK and Akt phosphorylation in murine primary macrophages.

Therefore, this readily available, inexpensive, and eco-friendly functional food may be a useful component in prophylactic strategies for regulating excessive inflammation in patients with COVID-19.

## Data Availability

The data supporting the findings in this study are available within the article.

## Conflicts of Interest

J. Takanari is an employee of Amino Up Co., Ltd. and any other authors declare that there are no conflicts of interest.

## Acknowledgments

This study was supported by a Grant-in-Aid for Scientific Research (C) (21K11472: K. Shirato) received from the Ministry of Education, Culture, Sports, Science and Technology, Japan. This study was also supported by the 36th Research Grant of the Meiji Yasuda Life Foundation of Health and Welfare (K. Shirato) and the 32nd Research Grant of the Nakatomi Foundation (K. Shirato).

## References

[1] The Novel Coronavirus Pneumonia Emergency Response Epidemiology Team, “The epidemiological characteristics of an outbreak of 2019 novel coronavirus diseases (COVID-19)–China, 2020,” China CDC Weekly, vol. 2, no. 8, pp. 113–122, 2020.

[2] Z. Wu and J. M. McGoogan, “Characteristics of and important lessons from the coronavirus disease 2019 (COVID-19) outbreak in China: summary of a report of 72 314 cases from the Chinese Center for Disease Control and Prevention,” Journal of the American Medical Association, vol. 323, no. 13, pp. 1239–1242, 2020.

[3] F. K. Ho, F. Petermann-Rocha, S. R. Gray et al., “Is older age associated with COVID-19 mortality in the absence of other risk factors? General population cohort study of 470,034 participants,” PLoS One, vol. 15, no. 11, Article ID e0241824, 2020.

[4] M. O’Driscoll, G. Ribeiro Dos Santos, L. Wang et al., “Age-specific mortality and immunity patterns of SARS-CoV-2,” Nature, vol. 590, no. 7844, pp. 140–145, 2021.

[5] M. R. Anderson, J. Geleris, D. R. Anderson et al., “Body mass index and risk for intubation or death in SARS-CoV-2 infection: a retrospective cohort study,” Annals of Internal Medicine, vol. 173, no. 10, pp. 782–790, 2020.

[6] L. Kompaniyets, A. B. Goodman, B. Belay et al., “Body mass index and risk for COVID-19-related hospitalization, intensive care unit admission, invasive mechanical ventilation, and death–United States, March–December 2020,” MMWR. Morbidity and Mortality Weekly Report, vol. 70, no. 10, pp. 355–361, 2021.

[7] S. Y. Tartof, L. Qian, V. Hong et al., “Obesity and mortality among patients diagnosed with COVID-19: results from an integrated health care organization,” Annals of Internal Medicine, vol. 173, no. 10, pp. 773–781, 2020.

[8] E. Barron, C. Bakhai, P. Kar et al., “Associations of type 1 and type 2 diabetes with COVID-19-related mortality in England: a whole-population study,” The Lancet. Diabetes & Endocrinology, vol. 8, no. 10, pp. 813–822, 2020.

[9] J. M. Dennis, B. A. Mateen, R. Sonabend et al., “Type 2 diabetes and COVID-19-related mortality in the critical care setting: a national cohort study in England, March– July 2020,” Diabetes Care, vol. 44, no. 1, pp. 50–57, 2021.

[10] N. Holman, P. Knighton, P. Kar et al., “Risk factors for COVID-19-related mortality in people with type 1 and type 2 diabetes in England: a population-based cohort study,” The Lancet. Diabetes & Endocrinology, vol. 8, no. 10, pp. 823–833, 2020.

[11] P. Mehta, D. F. McAuley, M. Brown, E. Sanchez, R. S. Tattersall, and J. J. Manson, “HLH across speciality collaboration, COVID-19: consider cytokine storm syndromes and immunosuppression,” Lancet, vol. 395, no. 10229, pp. 1033–1034, 2020.

[12] M. Merad and J. C. Martin, “Pathological inflammation in patients with COVID-19: a key role for monocytes and macrophages, Nature Reviews. Immunology, vol. 20, no. 6, pp. 355–362, 2020.

[13] G. Chen, D. Wu, W. Guo et al., “Clinical and immunological features of severe and moderate coronavirus disease 2019,” Journal of Clinical Investigation, vol. 130, no. 5, pp. 2620–2629, 2020.

[14] K. Shirato and T. Kizaki, “SARS-CoV-2 spike protein S1 subunit induces proinflammatory responses via Toll-like receptor 4 signaling in murine and human macrophages,” Heliyon, vol. 7, no. 2, Article ID e06187, 2021.

[15] E. Della-Torre, M. Lanzillotta, C. Campochiaro et al., “Respiratory impairment predicts response to IL-1 and IL-6 blockade in COVID-19 patients with severe pneumonia and hyper-inflammation,” Frontiers in Immunology, vol. 12, Article ID 675678, 2021.

[16] A. C. Gordon, P. R. Mouncey, F. Al-Beidh et al., “Interleukin-6 receptor antagonists in critically ill patients with covid-19,” New England Journal of Medicine, vol. 384, no. 16, pp. 1491–1502, 2021.

[17] T. Ito, K. Goto, J. Takanari et al., “Effects of enzyme-treated asparagus extract on heat shock protein 70, stress indices, and sleep in healthy adult men,” Journal of Nutritional Science and Vitaminology, vol. 60, no. 4, pp. 283–290, 2014.

[18] T. Ito, T. Maeda, K. Goto et al., “Enzyme-treated asparagus extract promotes expression of heat shock protein and exerts antistress effects,” Journal of Food Science, vol. 79, no. 3, pp. H413–H419, 2014.

[19] J. Takanari, J. Nakahigashi, A. Sato et al., “Effect of enzyme-treated asparagus extract (ETAS) on psychological stress in healthy individuals,” Journal of Nutritional Science and Vitaminology, vol. 62, no. 3, pp. 198–205, 2016.

[20] Z. Peng, S. Bedi, V. Mann et al., “Neuroprotective effects of Asparagus officinalis stem extract in transgenic mice overexpressing amyloid precursor protein,” Journal of Immunology Research, vol. 2021, Article ID 8121407, 2021.

[21] K. Shirato, J. Takanari, T. Koda et al., “A standardized extract of Asparagus officinalis stem prevents reduction in heat shock protein 70 expression in ultraviolet-B-irradiated normal human dermal fibroblasts: an in vitro study,” Environmental Health and Preventive Medicine, vol. 23, no. 1, Article ID 40, 2018.

[22] T. Ito, A. Sato, T. Ono et al., “Isolation, structural elucidation, and biological evaluation of a 5-hydroxymethyl-2-furfural derivative, asfural, from enzyme-treated asparagus extract,” Journal of Agricultural and Food Chemistry, vol. 61, no. 38, pp. 9155–9159, 2013.

[23] S. Inoue, J. Takanari, K. Abe, A. Nagayama, Y. Ikeya, and N. Kohda, “Isolation and structure determination of a heat shock protein inducer, asparagus -derived proline-containing 3-alkyldiketopiperazines (Asparaprolines), from a standardized extract of Asparagus officinalis stem,” Natural Product Communications, vol. 15, no. 3, pp. 1–7, 2020.

[24] K. Shirato, J. Takanari, J. Ogasawara et al., “Enzyme-treated asparagus extract attenuates hydrogen peroxide-induced matrix metalloproteinase-9 expression in murine skin fibroblast L929 cells,” Natural Product Communications, vol. 11, no. 5, pp. 677–680, 2016.

[25] K. Shirato, J. Takanari, T. Sakurai et al., “Enzyme-treated asparagus extract prevents hydrogen peroxide-induced pro-inflammatory responses by suppressing p65 nuclear translocation in skin L929 fibroblasts,” Natural Product Communications, vol. 11, no. 12, pp. 1883–1888, 2016.

[26] K. Shirato, T. Koda, J. Takanari et al., “Anti-inflammatory effect of ETAS®50 by inhibiting nuclear factor-κB p65 nuclear import in ultraviolet-B-irradiated normal human dermal fibroblasts,” Evidence-Based Complementary and Alternative Medicine, vol. 2018, Article ID 5072986, 2018.

[27] K. Shirato, T. Koda, J. Takanari et al., “ETAS®50 attenuates ultraviolet-B-induced interleukin-6 expression by suppressing Akt phosphorylation in normal human dermal fibroblasts,” Evidence-Based Complementary and Alternative Medicine, vol. 2018, Article ID 1547120, 2018.

[28] K. Shirato, K. Imaizumi, T. Sakurai, J. Ogasawara, H. Ohno, and T. Kizaki, “Regular voluntary exercise potentiates interleukin-1β and interleukin-18 secretion by increasing caspase-1 expression in murine macrophages,” Mediators of Inflammation, vol. 2017, Article ID 9290416, 2017.

[29] S. Kato, K. Shirato, K. Imaizumi et al., “Anticancer effects of phenoxazine derivatives combined with tumor necrosis factor-related apoptosis-inducing ligand on pancreatic cancer cell lines, KLM-1 and MIA-PaCa-2,” Oncology Reports, vol. 15, no. 4, pp. 843–848, 2006.

[30] E. L. Mills, B. Kelly, A. Logan et al., “Succinate dehydrogenase supports metabolic repurposing of mitochondria to drive inflammatory macrophages,” Cell, vol. 167, no. 2, pp. 457–470.e13, 2016.

[31] M. Pozzolini, S. Scarfi, U. Benatti, and M. Giovine, “Interference in MTT cell viability assay in activated macrophage cell line,” Analytical Biochemistry, vol. 313, no. 2, pp. 338–341, 2003.

[32] G. Arango Duque and A. Descoteaux, “Macrophage cytokines: involvement in immunity and infectious diseases,” Frontiers in Immunology, vol. 5, Article ID 491, 2014.

[33] K. V. Swanson, M. Deng, and J. P. Ting, “The NLRP3 inflammasome: molecular activation and regulation to therapeutics,” Nature Reviews. Immunology, vol. 19, no. 8, pp. 477–489, 2019.

[34] Y. S. Chang, B. H. Ko, J. C. Ju, H. H. Chang, S. H. Huang, and C. W. Lin, “SARS unique domain (SUD) of severe acute respiratory syndrome coronavirus induces NLRP3 inflammasome-dependent CXCL10-mediated pulmonary inflammation,” International Journal of Molecular Sciences, vol. 21, no. 9, Article ID 3179, 2020.

[35] P. C. Lara, D. Macías-Verde, and J. Burgos-Burgos, “Age-induced NLRP3 inflammasome over-activation increases lethality of SARS-CoV-2 pneumonia in elderly patients,” Aging and Disease, vol. 11, no. 4, pp. 756–762, 2020.

[36] S. J. Theobald, A. Simonis, T. Georgomanolis et al., “Long-lived macrophage reprogramming drives spike protein-mediated inflammasome activation in COVID-19,” EMBO Molecular Medicine, vol. 13, Article ID e14150, 2021.

[37] Z. Fan, L. Cai, Y. Wang, Q. Zhu, S. Wang, and B. Chen, “The acidic fraction of Isatidis Radix regulates inflammatory response in LPS-stimulated RAW264.7 macrophages through MAPKs and NF-κB Pathway,” Evidence-Based Complementary and Alternative Medicine, vol. 2021, Article ID 8879862, 2021.

[38] H. Y. Yu, K. S. Kim, Y. C. Lee, H. I. Moon, and J. H. Lee, “Oleifolioside A, a new active compound, attenuates LPS-stimulated iNOS and COX-2 expression through the downregulation of NF-κB and MAPK activities in RAW 264.7 macrophages,” Evidence-Based Complementary and Alternative Medicine, vol. 2012, Article ID 637512, 2012.

[39] G. R. Yu, S. J. Lee, D. H. Kim et al., “Literature-based drug repurposing in traditional Chinese medicine: reduced inflammatory M1 macrophage polarization by Jisil Haebaek Gyeji-Tang alleviates cardiovascular disease in vitro and ex vivo,” Evidence-Based Complementary and Alternative Medicine, vol. 2020, Article ID 8881683, 2020.

[40] H. Zhang, Q. Guo, Z. Liang et al., “Anti-inflammatory and antioxidant effects of Chaetoglobosin Vb in LPS-induced RAW264.7 cells: achieved via the MAPK and NF-κB signaling pathways,” Food and Chemical Toxicology, vol. 147, Article ID 111915, 2021.

[41] J. Guo, S. Tang, Y. Miao, L. Ge, J. Xu, and X. Zeng, “The anti-inflammatory effects of lignan glycosides from Cistanche tubulosa stems on LPS/IFN-γ-induced RAW264.7 macrophage cells via PI3K/AKT pathway,” Current Pharmaceutical Biotechnology, vol. 22, no. 10, pp. 1380–1391, 2021.

[42] M. A. Haque, I. Jantan, H. Harikrishnan, and W. Ahmad, “Standardized ethanol extract of Tinospora crispa upregulates pro-inflammatory mediators release in LPS-primed U937 human macrophages through stimulation of MAPK, NF-κB and PI3K-Akt signaling networks,” BMC Complementary Medicine and Therapies, vol. 20, no. 1, Article ID 245, 2020.

[43] R. S. Najjar, N. S. Akhavan, S. Pourafshar, B. H. Arjmandi, and R. G. Feresin, “Cornus officinalis var. koreana Kitam polyphenol extract decreases pro-inflammatory markers in lipopolysaccharide (LPS)-induced RAW 264.7 macrophages by reducing Akt phosphorylation,” Journal of Ethnopharmacology, vol. 270, Article ID 113734, 2021.

[44] S. Tanimura and K. Takeda, “ERK signalling as a regulator of cell motility,” Journal of Biochemistry, vol. 162, no. 3, pp. 145–154, 2017.

[45] J. S. Yu and W. Cui, “Proliferation, survival and metabolism: the role of PI3K/AKT/mTOR signalling in pluripotency and cell fate determination,” Development, vol. 143, no. 17, pp. 3050–3060, 2016.

[46] T. Ito, T. Ono, A. Sato et al., “Toxicological assessment of enzyme-treated asparagus extract in rat acute and subchronic oral toxicity studies and genotoxicity tests,” Regulatory Toxicology and Pharmacology, vol. 68, no. 2, pp. 240–249, 2014.

